# Proteolytic remodeling by Yme1 enables mitochondrial-derived compartment formation

**DOI:** 10.64898/2026.04.03.716366

**Authors:** Sai Sangeetha Balasubramaniam, Amy E. Curtis, Jonathan R. Friedman, Adam L. Hughes

## Abstract

Mitochondrial-derived compartments (MDCs) are remodeling domains that form from the outer mitochondrial membrane during metabolic and proteotoxic stress and selectively sequester hydrophobic membrane proteins. Although MDC formation depends on mitochondrial lipid composition and occurs at organelle contact sites, the molecular mechanisms that permit their biogenesis remain poorly defined. Here we identify the conserved inner mitochondrial membrane i-AAA protease Yme1 as a critical regulator of MDC formation. Loss of Yme1 blocks MDC biogenesis in response to multiple stressors, and this requirement depends on its proteolytic activity rather than secondary defects in mitochondrial morphology. Quantitative mitochondrial proteomics under MDC-inducing conditions revealed Yme1-dependent remodeling of lipid transfer proteins of the Ups family and components of the MICOS complex. Disruption of either pathway partially restores MDC formation in *yme1Δ* cells, while combined perturbation substantially bypasses the requirement for Yme1. Finally, Yme1 overexpression drives MDC formation in the absence of stress, although this activity remains constrained by metabolic conditions. Together, these findings support a model in which Yme1-dependent proteolysis relieves lipid- and MICOS-dependent constraints to permit MDC formation.

## INTRODUCTION

Mitochondrial plasticity enables cells to adapt to fluctuating physiological and nutrient conditions, thereby preserving cellular homeostasis (Cogliati *et al*, 2016; Bahat & Gross, 2019) This adaptability arises from coordinated functional and structural remodeling of both the outer and inner mitochondrial membranes (Pfanner *et al*, 2019; Colina-Tenorio *et al*, 2020). Mitochondrial membrane remodeling is therefore central to these adaptations, integrating changes in membrane architecture, lipid composition, and protein content in response to diverse stressors (Yang *et al*, 2022; Klecker & Westermann, 2021). These processes occur across a spectrum ranging from selective protein turnover mediated by mitochondrial proteases to large-scale degradation of entire organelles (Ng *et al*, 2021; Suomalainen & Nunnari, 2024). For example, mitophagy eliminates damaged mitochondria in response to severe or sustained stress, such as nutrient deprivation, hypoxia, mitochondrial poisons, or defects in protein import (Schuster & Okamoto, 2022; Abeliovich, 2023). In contrast, more selective remodeling pathways preserve mitochondrial integrity by removing or reorganizing specific components. These include mitochondrial-derived vesicles (MDVs), which form under steady-state conditions or during oxidative stress and encapsulate protein cargo from multiple mitochondrial subcompartments within membranes derived from the outer mitochondrial membrane or both mitochondrial membranes (Sugiura *et al*, 2014; Picca *et al*, 2023; König & McBride, 2024). Similarly, pathological perturbations, such as those observed during *Toxoplasma gondii* infection, remodel the outer mitochondrial membrane into large ring-shaped structures termed structures positive for the outer membrane (SPOTs), which incorporate specific OMM proteins while excluding intramitochondrial components (Li *et al*, 2022).

Among these pathways, a recently described mechanism of outer mitochondrial membrane remodeling is the mitochondrial-derived compartment (MDC) pathway (Hughes *et al*, 2016). MDCs are multilamellar structures that emerge from the outer mitochondrial membrane during metabolic and proteotoxic stress in budding yeast and mammalian cells (Schuler *et al*, 2020; Schuler *et al*, 2021; Wilson *et al*, 2024; Raghuram & Hughes, 2024). Under conditions such as elevated intracellular amino acid levels or disruptions in protein homeostasis, the outer mitochondrial membrane undergoes successive rounds of elongation and invagination to generate MDCs (Schuler *et al*, 2020; Wilson *et al*, 2024). Pharmacological perturbations that elevate intracellular amino acid pools in yeast, including inhibition of vacuolar amino acid storage with concanamycin A (ConcA) or translational inhibition with rapamycin (Rap) or cycloheximide (CHX), robustly induce MDC formation (Hughes *et al*, 2016; Schuler *et al*, 2021).

Upon formation, MDCs selectively sequester the mitochondrial import receptor Tom70 together with a defined subset of outer mitochondrial membrane proteins, while excluding proteins from other mitochondrial subcompartments (Hughes *et al*, 2016; Schuler *et al*, 2021; Wilson *et al*, 2024). These structures therefore represent specialized mitochondrial remodeling domains that capture and compartmentalize hydrophobic outer membrane cargo during metabolic and proteotoxic stress (Wilson *et al*, 2024; Raghuram & Hughes, 2024). MDCs arise through successive rounds of outer mitochondrial membrane elongation and invagination, generating multilamellar compartments composed exclusively of outer membrane (Wilson *et al*, 2024). Previous work, including an unbiased genome-wide genetic screen for regulators of MDC formation, identified mitochondrial lipid homeostasis as a critical determinant of this pathway (Xiao *et al*, 2024). In particular, mitochondrial lipid transfer proteins of the Ups family and components of the ER–mitochondria encounter structure (ERMES) were found to be required for MDC biogenesis, indicating that mitochondrial lipid composition and membrane organization play central roles in generating these compartments (English *et al*, 2020; Xiao *et al*, 2024). Despite these advances, the molecular mechanisms that drive MDC formation remain poorly understood, and it is still unclear how metabolic or proteotoxic stress is translated into the lipid and membrane remodeling events that produce these cargo-sequestering structures.

Among the candidates identified in this previous screen for MDC regulators was the conserved inner mitochondrial membrane i-AAA protease Yme1, an inner mitochondrial membrane (IMM)-localized mitochondrial protease known to regulate membrane protein turnover and mitochondrial membrane organization (Weber *et al*, 1996; Leonhard *et al*, 1999; Deshwal *et al*, 2020; Kan *et al*, 2024). Given the emerging importance of mitochondrial lipid composition and membrane architecture in MDC formation, we therefore investigated whether Yme1 contributes to MDC biogenesis.

In this study, we find that Yme1 proteolytic activity is required for MDC formation in response to multiple metabolic stress conditions that induce MDCs. Quantitative mitochondrial proteomics under MDC-inducing conditions revealed Yme1-dependent remodeling of lipid transfer proteins of the Ups family and components of the MICOS complex. Genetic perturbation of these pathways partially restores MDC formation in *yme1Δ* cells, while combined disruption substantially bypasses the requirement for Yme1. Together, these findings identify Yme1 proteolysis as a key regulator of MDC biogenesis and support a model in which Yme1-dependent remodeling of mitochondrial lipid transfer and membrane-organizing complexes relieves constraints that otherwise limit MDC formation.

## RESULTS AND DISCUSSION

### Proteolytic activity of Yme1 is required for MDC formation

To investigate the potential role of Yme1 in the MDC pathway, we examined the impact of *YME1* deletion on MDC formation across several well-characterized MDC inducers. MDCs were quantified using a fluorescence microscopy assay in which MDCs appear as discrete Tom70-positive puncta that are enriched for outer mitochondrial membrane proteins while excluding inner mitochondrial membrane markers such as Tim50, thereby distinguishing them from the contiguous mitochondrial network (Hughes *et al*, 2016; Fig. 1A). Consistent with the preliminary results from our previous MDC screen, we found that deletion of *YME1* blocked MDC formation in response to a 2-hour exposure to Rap, ConcA, and CHX (Fig. 1A-B, Fig. S1A).

**Figure 1:**
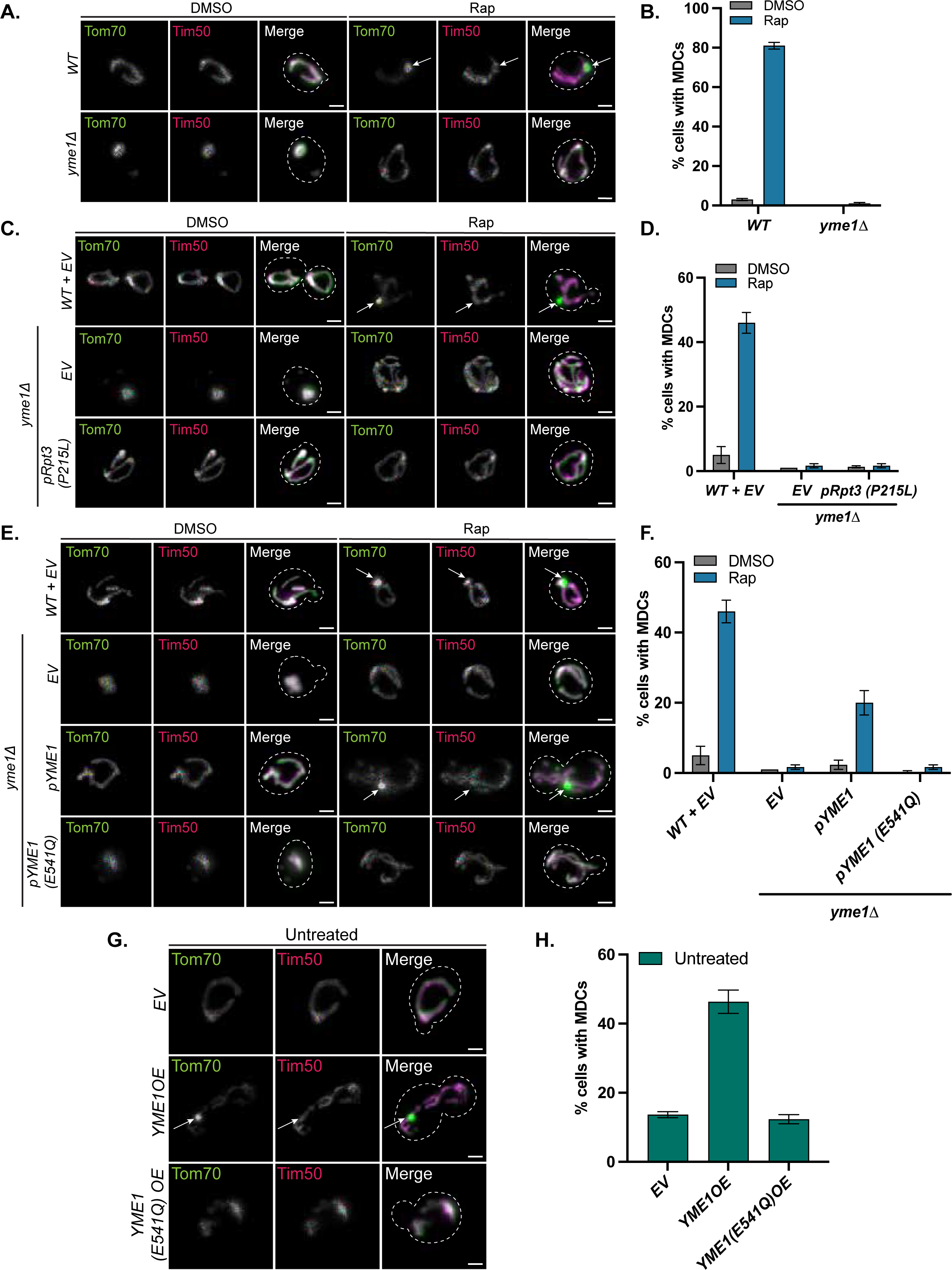
Proteolytic activity of Yme1 is required for MDC formation (A) Widefield images of wild-type and *yme1Δ* cells expressing Tom70-GFP and Tim50-mCherry treated with DMSO or Rap for 2 h. White arrows denote MDCs. Scale bar = 2 µm. (B) Quantification of (A) showing the percentage of cells with MDCs. Error bars represent the mean ± SE of three replicates, *n* ≥ 100 cells per replicate. (C) Widefield images of wild-type and *yme1Δ* cells endogenously tagged with Tom70-GFP and Tim50-mCherry expressing either *EV* or *pRS413-Rpt3 (P215L)* treated with DMSO or Rap for 2 h. White arrows denote MDCs. Scale bar = 2 µm. (D) Quantification of (C) showing the percentage of cells with MDCs. Error bars represent the mean ± SE of three replicates, *n* ≥ 100 cells per replicate. (E) Widefield images of wild-type and *yme1Δ* cells endogenously tagged with Tom70-GFP and Tim50-mCherry expressing either *EV*, *pRS413-Yme1*, or *pRS413-Yme1 (E541Q)* treated with DMSO or Rap for 2 h. White arrows denote MDCs. Scale bar = 2 µm. (F) Quantification of (E) showing the percentage of cells with MDCs. Error bars represent the mean ± SE of three replicates, *n* ≥ 100 cells per replicate. (G) Widefield images of yeast cells tagged with Tom70-GFP and Tim50-mCherry, constitutively overexpressing *YME1* (*YME1 OE*), the protease-dead mutant, *YME1(E541Q)*, or an empty vector (*EV*) control. White arrows denote MDCs. Scale bar = 2 µm. (H) Quantification of (G) showing the percentage of cells with MDCs. Error bars represent the mean ± SE of three replicates, *n* ≥ 100 cells per replicate.

Cells lacking Yme1 exhibit aberrant mitochondrial morphology, a high rate of mitochondrial DNA escape, and growth defects on nonfermentable media (Thorsness & Fox, 1993; Weber *et al*, 1996). Because loss of *YME1* disrupts mitochondrial morphology, we asked whether the MDC defect in *yme1Δ* cells arises as a secondary consequence of these structural alterations. To test this, we examined whether restoring mitochondrial morphology rescues MDC formation in *yme1Δ* mutants. Expression of a mutant form of the proteasome component Rpt3 (P215L) has been reported to suppress certain growth defects and alleviate mitochondrial morphological abnormalities in cells lacking *YME1* (Campbell *et al* 1994; Francis & Thorsness, 2011). Consistent with previous observations, we found that strains expressing *pRPT3(P215L)* exhibited substantial recovery of mitochondrial morphology but still failed to form MDCs (Fig. 1C-D, Fig. S1B-D). Moreover, Rap treatment also restored mitochondrial morphology in *yme1Δ* mutants through an unknown mechanism despite MDCs remaining absent (Fig. S1E). To determine whether loss of mtDNA contributes to the MDC defect in *yme1Δ* mutants, we examined MDC formation in wild-type cells lacking mitochondrial DNA (rho⁰). We found that rho⁰ cells exhibited MDC formation even in the absence of stress, indicating that mtDNA is not required for MDC induction (Fig. S1F-G). Together, these results indicate that neither altered mitochondrial morphology nor loss of mtDNA accounts for the MDC defect observed in *yme1Δ* cells.

To test whether the protease activity of Yme1 is required for MDC formation, we generated *yme1Δ* strains ectopically expressing either wild-type Yme1 or a catalytically inactive mutant (*yme1^E541Q^*) (Leonhard *et al*, 1999). Expression of wild-type Yme1 moderately restored MDC formation, whereas the protease mutant failed to rescue MDC formation in cells lacking *YME1* (Fig. 1E-F, Fig. S1H). Finally, we found that overexpression of wild-type Yme1 from a strong, constitutive GPD promoter, but not the catalytically inactive mutant, was sufficient to trigger MDC formation in in the absence of stress (Fig. 1G-H). These results indicate that elevated Yme1 levels can drive MDC formation in a manner dependent on its proteolytic activity Collectively, these data establish that Yme1 proteolytic activity is critical for MDC biogenesis.

### MDC Induction Triggers Yme1-Dependent Changes in the Mitochondrial Proteome

Given that the proteolytic activity of Yme1 is required for MDC biogenesis, we hypothesized that Yme1-dependent remodeling of mitochondrial proteins may be important for regulation of this pathway. To identify potential candidates, we performed TMT-based quantitative proteomics on mitochondria isolated from wild-type and *yme1Δ* cells in the absence and presence of a two-hour Rap treatment. Proteomic profiling identified more than 4,000 proteins and revealed extensive remodeling of the mitochondrial proteome upon MDC induction in a Yme1-dependent manner. Quality control analyses, including principal component analysis and normalization across samples, confirmed robust separation of the experimental groups and successful dataset normalization (Fig. S2A-B).

To identify broader biological patterns within the dataset, we grouped proteins based on shared changes in abundance across conditions. Hierarchical clustering identified distinct functional groups encompassing mitochondrial protein import, membrane organization, metabolism, and lipid homeostasis, several of which have previously been linked to MDC biogenesis (Fig. 2A). Thus, MDC induction is accompanied by broad remodeling of mitochondrial pathways, and loss of Yme1 alters multiple protein groups likely relevant to MDC formation.

**Figure 2:**
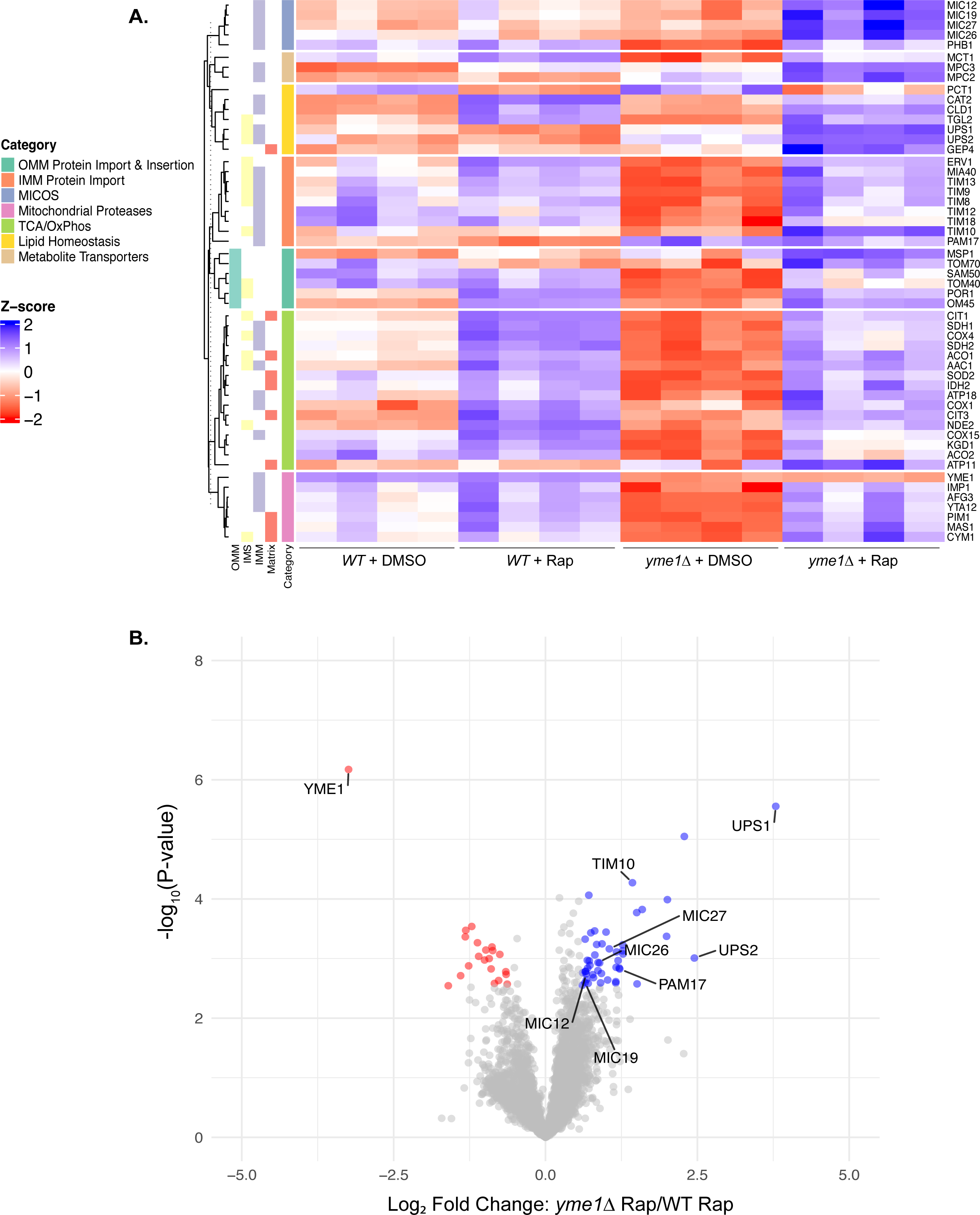
MDC Inducer Rapamycin triggers Yme1-dependent changes in the mitochondrial proteome (A) Clustered heatmap showing *Z*-scores of select mitochondrial proteins in wild-type and *yme1Δ* cells treated with DMSO or Rap for 2 h (n = 4), with corresponding mitochondrial subcategories indicated on the left. Proteins were categorized by GO Biological Processes (GO: BP) and manual annotation. Blue indicates upregulated proteins; red indicates downregulated proteins. (B) Volcano plot analysis of differentially expressed proteins in *yme1Δ* cells compared with wild-type cells following Rap treatment for 2 h (n = 4). Mitochondrial proteins enriched in *yme1Δ* cells are highlighted in blue and represent potential Yme1 substrates; red indicates downregulated proteins.

To identify the proteins most strongly affected by loss of Yme1 under MDC-inducing conditions that may be good candidates for MDC regulators, we next compared the Rap-treated wild-type and *yme1Δ* mitochondrial proteomes by volcano plot analysis (Fig. 2B). Among the most significantly elevated proteins in *yme1Δ* cells were Ups1 and Ups2, members of the conserved Ups/PRELID family of mitochondrial lipid transfer proteins that shuttle phospholipid precursors between mitochondrial membranes (Potting *et al*, 2010; Tamura *et al*, 2012; Connerth *et al*, 2012; Aaltonen *et al*, 2016; Tatsuta & Langer, 2017; MacVicar *et al*, 2019; Perea *et al*, 2023). In addition, several components of the MICOS complex (mitochondrial contact site and cristae organizing system) were also elevated (Schreiner *et al*, 2012; Li *et al*, 2016). MICOS is a conserved multiprotein complex located at crista junctions that maintains inner mitochondrial membrane architecture and coordinates membrane contact sites between mitochondrial subcompartments (Aaltonen *et al*, 2016; Cogliati *et al*, 2016; Huynen *et al*, 2016; Mukherjee *et al*, 2021). The elevated protein dataset also included proteins involved in mitochondrial protein import, including members of the small TIM chaperone complexes, which facilitate the transfer of hydrophobic precursor proteins across the intermembrane space (Koehler *et al*, 1998; Koehler, 1998; Baker *et al*, 2012; Spiller *et al*, 2015), and Pam17, a regulatory component of the mitochondrial inner membrane import motor that assists in translocation of proteins into the matrix (Van Der Laan *et al*, 2005; Schiller, 2009). Notably, many of these proteins reside within mitochondrial subcompartments accessible to Yme1 proteolysis, including the IMM and intermembrane space (IMS), and several are well-established Yme1-regulated substrates, including the Ups proteins. Together, these results show that loss of Yme1 alters the abundance of proteins across multiple mitochondrial pathways under MDC-inducing conditions, providing a pool of candidate regulators for further investigation.

### Yme1 regulation of Ups2 contributes to MDC formation

To determine whether proteins elevated in *yme1Δ* cells might represent Yme1-regulated factors that influence MDC biogenesis, we next examined candidates emerging from the proteomic analysis. Because MDC formation is completely blocked in *yme1Δ* cells while several proteins accumulate under these conditions, we reasoned that elevation of these proteins might inhibit MDC formation or that their ongoing activity may normally act as a constraint on MDC biogenesis. Among the strongest candidates were the mitochondrial lipid transfer proteins Ups1 and Ups2, which were significantly elevated in Rap-treated *yme1Δ* cells (Fig. 2B). The Ups proteins regulate mitochondrial lipid composition by shuttling phospholipid precursors from the outer mitochondrial membrane to the inner mitochondrial membrane (Tamura *et al*, 2012; Connerth *et al*, 2012; Aaltonen *et al*, 2016). Ups1 transfers phosphatidic acid (PA) for downstream cardiolipin synthesis, whereas Ups2 transports phosphatidylserine (PS) to facilitate phosphatidylethanolamine (PE) production. Previous work has shown that Yme1 mediates Ups2 turnover, while Ups1 degradation can be regulated by both Yme1 and the metallopeptidase Atp23 (Potting *et al*, 2010; Tatsuta & Langer, 2017).

To test whether the changes observed in our proteomic analysis reflect Yme1-dependent regulation of Ups proteins, we examined Ups1 and Ups2 protein levels by immunoblotting. Consistent with the proteomics dataset, Ups1 and Ups2 steady-state levels declined in wild-type cells following Rap treatment but accumulated and remained elevated in *yme1Δ* mutants under the same conditions (Fig. 3A-D). These results confirm that Ups proteins are regulated by Yme1 and that their abundance decreases under MDC-inducing conditions in a Yme1-dependent manner.

**Figure 3:**
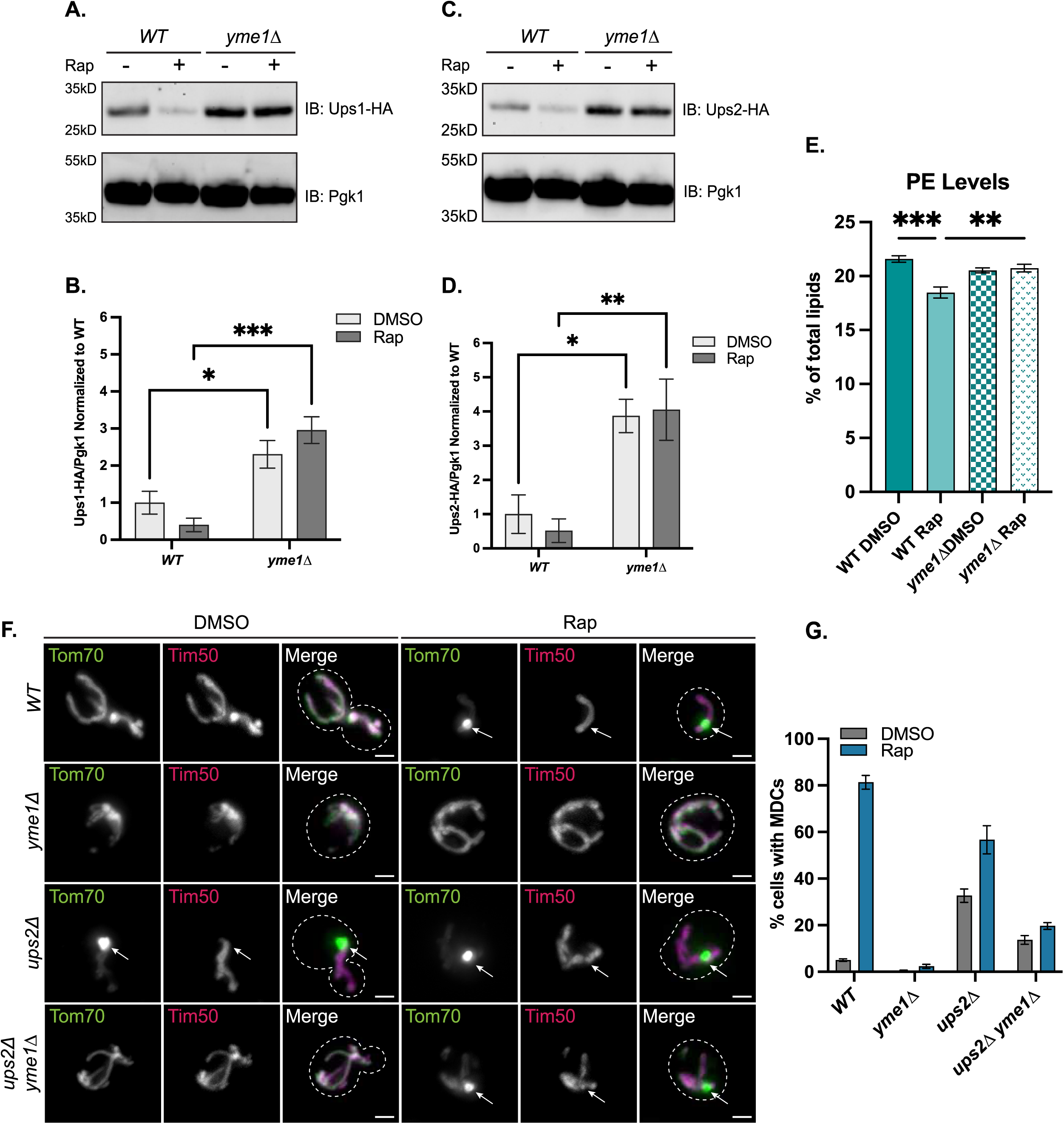
Yme1 regulation of Ups2 contributes to MDC formation (A) Immunoblot analysis of whole-cell lysates from yeast expressing endogenously tagged Ups1-HA, treated with DMSO or Rap for 2 h. Blots were probed with anti-HA; Pgk1 served as a loading control. Representative of n = 3 independent experiments. (B) Quantification of (A) wherein Ups1-HA levels were normalized to the corresponding Pgk1 signal and then normalized to *EV* cells treated with DMSO (n = 3). Statistical significance was determined by two-way ANOVA (*p ≤ 0.05, ***p ≤ 0.001). (C) Immunoblot analysis of whole-cell lysates from yeast expressing endogenously tagged Ups2-HA, treated with DMSO or Rap for 2 h. Blots were probed with anti-HA; Pgk1 served as a loading control. Representative of n = 3 independent experiments. (D) Quantification of (C) wherein Ups2-HA levels were normalized to the corresponding Pgk1 signal and then normalized to *EV* cells treated with DMSO (n = 3). Statistical significance was determined by two-way ANOVA (*p ≤ 0.05, **p ≤ 0.01). (E) Relative whole-cell PE levels in wild-type and *yme1Δ* cells treated with DMSO or Rap for 2 h. PE levels are normalized to total lipids. Error bars represent the mean ± SE of four biological replicates. Statistical significance was determined by two-way ANOVA with Holm-Šídák post hoc test comparing each condition to its corresponding DMSO control. (**p ≤ 0.01, ***p ≤ 0.001). (F) Widefield images of wild-type and indicated mutants expressing Tom70-GFP and Tim50-mCherry treated with DMSO or Rap for 2 h. White arrows denote MDCs. Scale bar = 2 µm. (G) Quantification of (F) showing the percentage of cells with MDCs. Error bars represent the mean ± SE of three replicates, *n* ≥ 100 cells per replicate.

Previous work from our laboratory demonstrated that mitochondrial lipid remodeling is important for MDC biogenesis. Specifically, we showed that metabolic stress triggers changes in mitochondrial lipid composition, including a reduction in phosphatidylethanolamine (PE) that is required for MDC formation (Xiao *et al*, 2024). Consistent with this, we previously found that deletion of *UPS2* leads to constitutive MDC formation. How these lipid changes are triggered during MDC induction has remained unclear. The observation that Ups2 accumulates in *yme1Δ* mutants suggested that Yme1 might regulate MDC formation by altering Ups2 abundance. Consistent with this idea, we found that the Rap-induced decline in whole-cell PE previously associated with MDC induction was blunted in *yme1Δ* cells (Fig. 3E). Although the change in PE measured at the whole-cell level is modest, likely due to contributions from non-mitochondrial lipid pools, our previous work demonstrated that similarly small changes in total cellular PE are sufficient to regulate MDC formation (Xiao et al, 2024, Fig. 3E). We next tested whether removal of Ups2 could bypass the requirement for Yme1 in the MDC pathway. In *yme1Δ ups2Δ* double mutants, MDC formation was partially restored. Approximately 14% of untreated double mutant cells formed MDCs, and Rap treatment modestly increased this to ∼20% (Fig. 3F–G). Similar trends were observed following treatment with ConcA and CHX, with MDC formation reaching ∼24% and ∼20%, respectively (Fig. S3A). Notably, MDC formation in the double mutant remained lower than in *ups2Δ* cells alone. These results suggest that while accumulation of Ups2 contributes to the MDC defect in *yme1Δ* cells, removal of Ups2 is not sufficient to fully restore MDC biogenesis in the absence of Yme1. Together, these findings indicate that Yme1 likely promotes MDC formation in part through regulation of mitochondrial lipid transfer pathways and control of PE abundance, but that additional Yme1-regulated factors also contribute to this process.

### MICOS complex contributes to Yme1-dependent regulation of MDC formation

The partial rescue of MDC formation in *ups2Δ yme1Δ* mutants suggested that Yme1 regulates the abundance of additional factors that influence MDC biogenesis. Re-examining the proteomics dataset revealed that several components of the MICOS complex—including Mic12, Mic19, Mic26, and Mic27—were significantly elevated in Rap-treated *yme1Δ* cells (Fig. 4A), suggesting that Yme1 may also regulate additional pathways linked to mitochondrial membrane organization, including the MICOS complex, during MDC induction.

**Figure 4:**
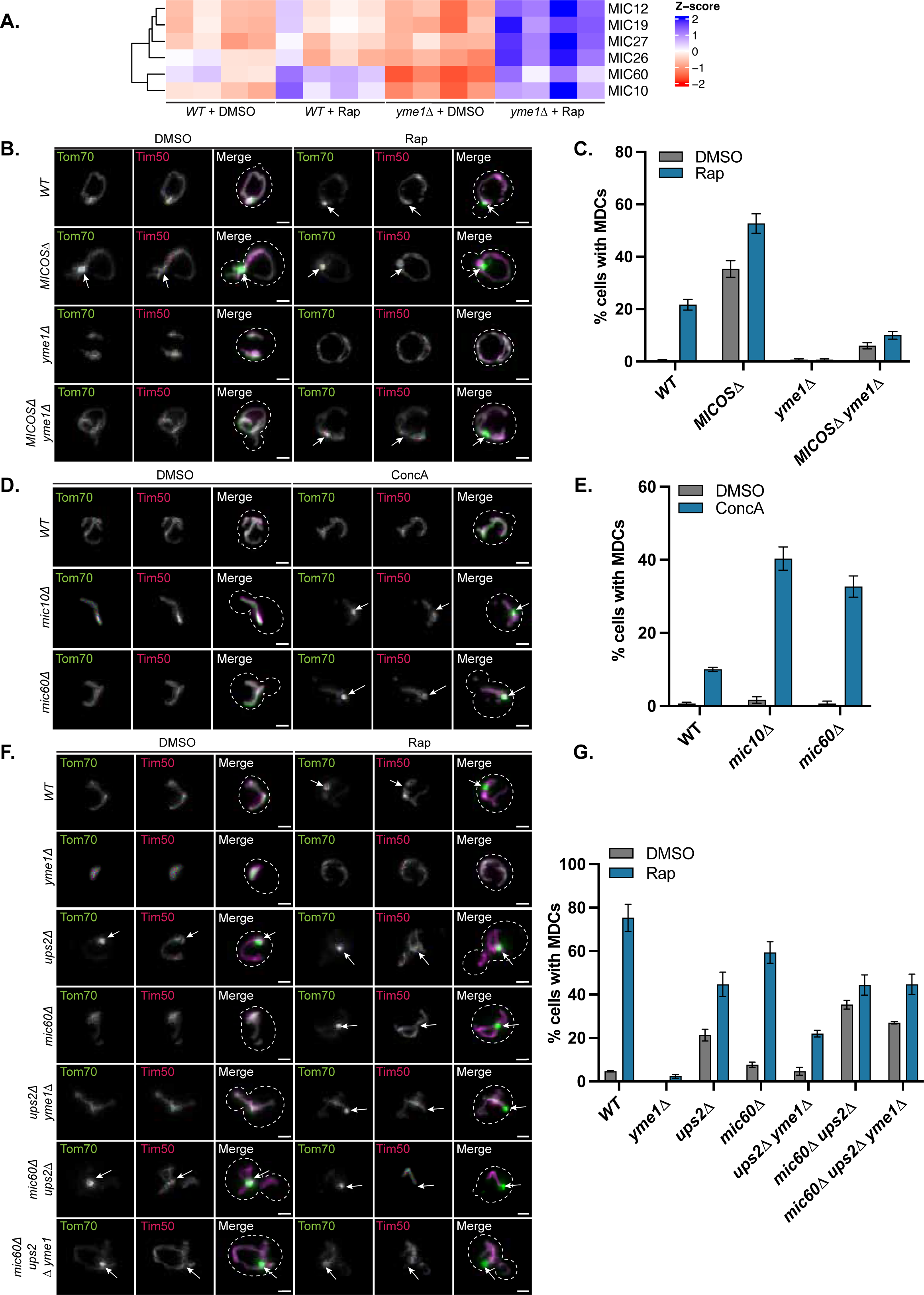
MICOS complex contributes to Yme1-dependent regulation of MDC formation (A) Heatmap showing *Z*-scores of MICOS proteins in wild-type and *yme1Δ* cells treated with DMSO or Rap for 2 h (n = 4). Blue indicates upregulated proteins; red indicates downregulated proteins. (B) Widefield images of wild-type and indicated mutants expressing Tom70-GFP and Tim50-mCherry treated with DMSO or Rap for 2 h. White arrows denote MDCs. Scale bar = 2 µm. (C) Quantification of (B) showing the percentage of cells with MDCs. Error bars represent the mean ± SE of three replicates, *n* ≥ 100 cells per replicate. (D) Widefield images of wild-type and indicated mutants expressing Tom70-GFP and Tim50-mCherry, grown in media containing low amino acids and treated with DMSO or ConcA for 2 h. White arrows denote MDCs. Scale bar = 2 µm. (E) Quantification of (D) showing the percentage of cells with MDCs. Error bars represent the mean ± SE of three replicates, *n* ≥ 100 cells per replicate. (F) Widefield images of wild-type and indicated mutants expressing Tom70-GFP and Tim50-mCherry treated with DMSO or Rap for 2 h. White arrows denote MDCs. Scale bar = 2 µm. (G) Quantification of (F) showing the percentage of cells with MDCs. Error bars represent the mean ± SE of three replicates, *n* ≥ 100 cells per replicate.

The MICOS complex is a conserved multiprotein complex that resides at cristae junctions and maintains the structural organization of the inner mitochondrial membrane (Friedman *et al*, 2015; Huynen *et al*, 2016; Mukherjee *et al*, 2021). In yeast, MICOS consists of six subunits organized into two functional modules: a Mic60–Mic19 subcomplex and a Mic10-containing subcomplex comprising Mic10, Mic12, Mic26, and Mic27 (Friedman *et al*, 2015; Huynen *et al*, 2016;Bohnert *et al*, 2015). Together with ATP synthase dimers, MICOS regulates cristae architecture and also helps maintain connectivity between the inner and outer mitochondrial membranes (Ott *et al*, 2012; Körner *et al*, 2012; Tang *et al*, 2020). Because multiple MICOS components accumulated in *yme1Δ* cells upon Rap treatment, we asked whether disruption of this entire complex influences MDC formation. As the complete MICOS deletion strain lacking all MICOS subunits was generated and characterized previously in the W303 genetic background (Friedman *et al*, 2015), we first verified that MDC formation occurs in this strain background, given that all prior experiments were conducted in the S288C (BY) background. Indeed, Rap treatment induced MDC formation in wild-type W303 cells, although at lower levels (∼20%) than typically observed in BY strains (∼70%), establishing a lower baseline for this background (Fig. 4B–C). Strikingly, deletion of the MICOS complex resulted in constitutive MDC formation, with ∼35% of untreated *MICOSΔ* cells forming MDCs. Rap treatment further increased MDC formation to nearly 50% (Fig. 4B–C). Similar increases were observed following ConcA and CHX treatment (Fig. S3B). These results suggest that disruption of the MICOS complex stimulates MDC biogenesis and that this protein complex normally acts as a constraint on MDC formation.

We next tested whether MICOS deletion could restore MDC formation in *yme1Δ* mutants. While deletion of *YME1* alone blocked MDC formation in W303 cells, MDC formation was partially restored in *MICOSΔ yme1Δ* mutants. Following Rap treatment, approximately 10% of these cells formed MDCs—about half the level observed in wild-type W303 cells (∼20%) (Fig. 4B–C). Similar partial restoration was observed following ConcA and CHX treatment (Fig. S3B). These results indicate that altered MICOS abundance contributes to the MDC defect in *yme1Δ* cells but that loss of this complex is not sufficient to fully bypass the requirement for Yme1, similar to the partial bypass effects overserved with loss of Ups2 in strains lacking Yme1.

### Combined loss of MICOS components and Ups2 bypasses Yme1 in MDC formation

Because deletion of *UPS2* or disruption of the MICOS complex each partially restored MDC formation in *yme1Δ* cells, we next asked whether combining these perturbations could further bypass the requirement for Yme1. Before testing this interaction, we first tested whether deletion of *UPS2* promotes constitutive MDC formation in the W303 background as it does in the BY background. Indeed, ∼30% of untreated ups2Δ W303 cells formed MDCs, with Rap treatment modestly increasing MDC formation to ∼40% (Fig. S3C-D). Next, we tested whether simultaneous disruption of MICOS and Ups2 could enhance MDC formation. Indeed, *ups2Δ MICOSΔ* mutants exhibited elevated constitutive MDC formation (∼40%), higher than either single mutant, suggesting that lipid transfer (UPS proteins) and mitochondrial architecture (MICOS) represent independent constraints on MDC formation (Fig. S3C-D).

To determine whether loss of both of these pathways could fully bypass Yme1, we attempted to construct a *MICOSΔ ups2Δ yme1Δ* triple mutant. However, this strain was not viable, preventing direct analysis of complete MICOS disruption in this context. To overcome this limitation, we examined individual MICOS subunits to identify components with the strongest influence on MDC formation. Deletion strains were generated in the BY background and tested for MDC formation. Unlike the full MICOS deletion, under nutrient-rich conditions, deletion of individual MICOS subunits did not constitutively activate MDC formation nor did it significantly alter MDC formation upon ConcA, Rap, or CHX treatment compared to wild-type cells (Fig. S3E). However, when MDC formation was examined under metabolically restrictive conditions (low amino acid media), which normally block ConcA-induced MDC formation, deletion of specific MICOS components restored MDC formation. Under these conditions, *mic10Δ* mutants displayed MDC formation in ∼40% of ConcA-treated cells and mic60Δ mutants in ∼30%, compared to ∼10% in wild-type (Fig. 4D–E). Deletion of additional MICOS subunits (mic27Δ, mic12Δ, and mic19Δ) also increased MDC formation to a lesser extent, while mic26Δ had little effect (Fig. S4A–B), indicating that loss of MICOS components variably sensitizes cells to MDC formation.

Based on these results, we generated triple-mutant strains combining *ups2Δ* and *yme1Δ* with deletion of either *MIC10* or *MIC60*. The *mic60Δ ups2Δ yme1Δ* strain showed substantially restored MDC formation, with ∼27% of untreated cells forming MDCs and ∼45% forming MDCs following Rap treatment (Fig. 4F-G). These results indicate that simultaneous disruption of Ups2 and Mic60 can largely bypass the requirement for Yme1. Notably, a *mic60Δ yme1Δ* double mutant could not be generated due to synthetic lethality in the BY background, highlighting a strong functional interaction between these pathways (Fig. S4C). The *mic10Δ ups2Δ yme1Δ* strain exhibited partial rescue of basal MDC formation (∼24%) but did not respond strongly to MDC inducers (Fig. S4D-E). Together, these findings indicate that multiple mitochondrial constraints—including lipid composition and MICOS—must be altered to permit MDC formation, and that Yme1 promotes MDC biogenesis by coordinately regulating these pathways.

### Yme1-dependent MDC formation is gated by metabolic cues

Having established that Yme1 controls MDC formation by regulating lipid transfer proteins and MICOS, we next asked how this activity is controlled and whether increasing Yme1 levels is sufficient to drive MDC biogenesis. As shown in Figure 1G–H, overexpression of Yme1 triggered MDC formation in a large fraction of untreated cells, and this effect required Yme1 proteolytic activity, as a catalytically inactive mutant failed to induce MDCs. Importantly, several of the Yme1-regulated factors identified in our proteomic analysis—including Ups proteins and MICOS components—are altered in abundance even at steady state in the absence of Yme1 and become further altered during MDC-inducing conditions such as Rap treatment. These observations suggest that Yme1-dependent regulation of mitochondrial proteins occurs constitutively, but is amplified during stress, resulting in greater remodeling of lipid transfer pathways and MICOS. Consistent with this idea, Yme1 overexpression-driven MDC formation remained dependent on metabolic context. In contrast to overexpressing Yme1 in nutrient rich medium (Fig. 1G-H), overexpressing Yme1 in minimal media lacking amino acids did not stimulate MDC formation (Fig. 5A–B). Thus, even when Yme1 levels are elevated, MDC biogenesis requires a permissive metabolic environment.

**Figure 5:**
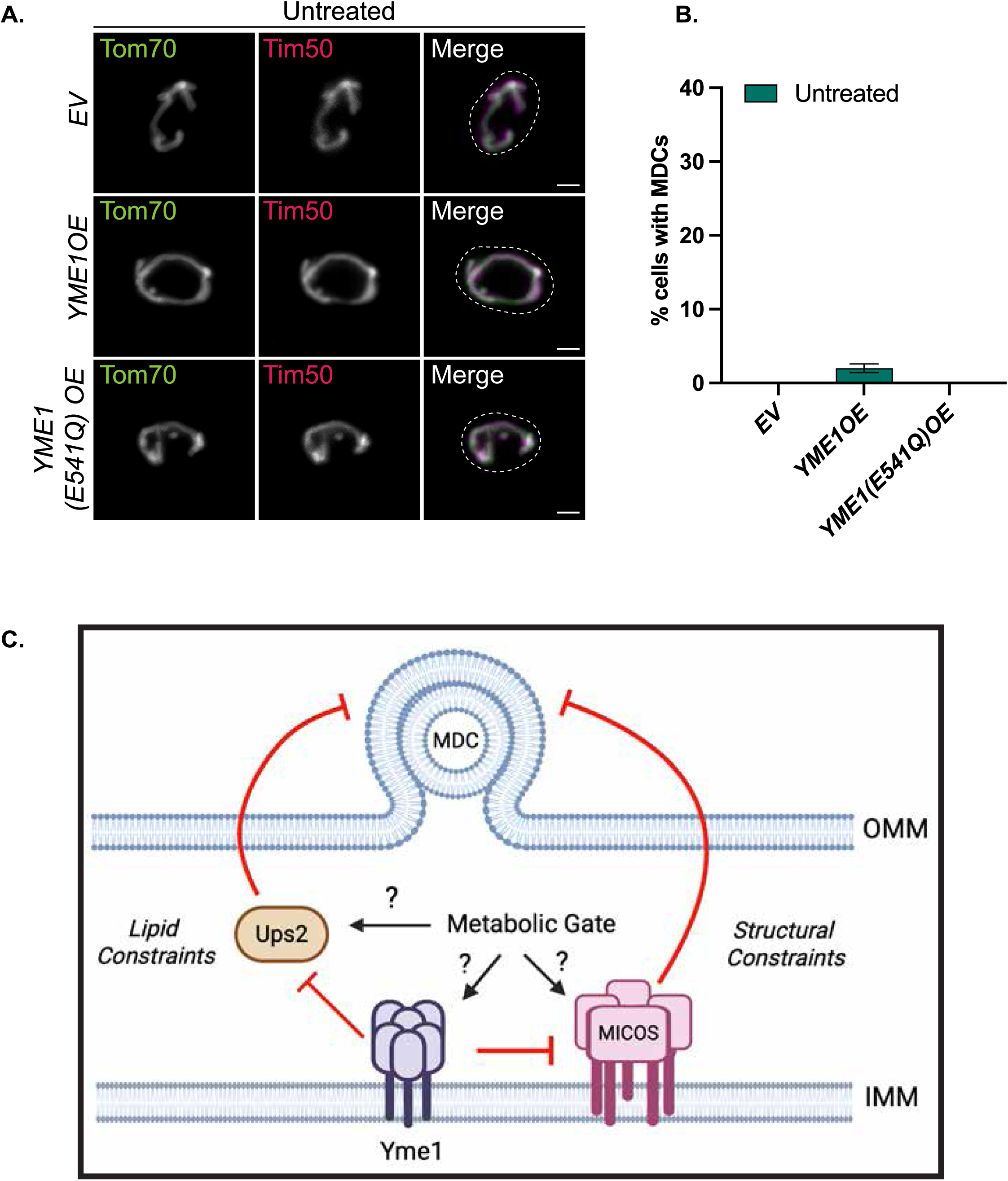
Yme1-dependent MDC formation is gated by metabolic cues (A) Widefield images of yeast cells tagged with Tom70-GFP and Tim50-mCherry, constitutively overexpressing *YME1* (*YME1 OE*), the protease-dead mutant, *YME1(E541Q)*, or an empty vector (*EV*) control grown in minimal media that lacks amino acids. Scale bar = 2 µm. (B) Quantification of (A) showing the percentage of cells with MDCs. Error bars represent the mean ± SE of three replicates, *n* ≥ 100 cells per replicate. (C) Model of Yme1-mediated MDC biogenesis: Upon treatment with MDC stimulating stressors, Yme1 degrades Ups2, altering lipids, while also modulating the MICOS complex to possibly alter organizational or structural constraints that normally inhibit MDC formation (Created in BioRender. Bala, S. (2026) https://BioRender.com/8srjgaz)

Together with the genetic and proteomic observations described above, these findings support a model in which Yme1 promotes MDC biogenesis by progressively relieving lipid and membrane organizational constraints imposed by the Ups pathway and MICOS. Under basal conditions, Yme1 activity likely contributes to ongoing mitochondrial remodeling but is insufficient to trigger widespread MDC formation. During metabolic stress, however, this remodeling is amplified—either through increased Yme1 activity or through changes in substrate susceptibility that enhance Yme1-dependent turnover—thereby permitting MDC biogenesis. The inability of Yme1 overexpression to induce MDCs under restrictive metabolic conditions further indicates that additional metabolic inputs gate pathway activation. This model, summarized below and provided Figure 5C, places Yme1 as a central regulator that removes constraints on MDC formation, while metabolic signals determine when this remodeling is sufficient to drive the pathway.

### Summary and model for Yme1-dependent regulation of mitochondrial-derived compartment biogenesis

Mitochondrial membranes dynamically remodel in response to metabolic shifts and cellular stress. In previous work, we identified the mitochondrial-derived compartment (MDC) pathway as a mechanism of outer mitochondrial membrane remodeling that selectively sequesters hydrophobic OMM proteins during metabolic and proteotoxic stress. MDC formation was previously shown to depend on metabolic signals that alter mitochondrial lipid composition, particularly reductions in phosphatidylethanolamine (PE). However, the molecular mechanisms that translate these metabolic signals into the membrane remodeling events required for MDC formation have remained unclear.

In this study, we identify the conserved mitochondrial i-AAA protease Yme1 as a central regulator of MDC biogenesis. Through genetic, proteomic, and biochemical approaches, we find that Yme1 promotes MDC formation by remodeling multiple mitochondrial pathways that normally act as constraints on OMM remodeling. Our data implicate the Ups lipid transfer pathway and the MICOS complex, two systems that regulate mitochondrial lipid homeostasis and membrane organization. Disruption of either pathway partially restores MDC formation in *yme1Δ* cells, whereas combined perturbation of both pathways largely bypasses the requirement for Yme1. These findings imply that MDC formation may require coordinated remodeling of both mitochondrial lipid composition and mitochondrial membrane organization.

How these pathways constrain MDC formation remains an important open question. One possibility is that Yme1-mediated turnover of Ups proteins alters mitochondrial lipid composition and/or redistributes lipids between the IMM and OMM, including local PE levels, thereby promoting membrane curvature or generating OMM membrane domains permissive for MDC formation (Agrawal & Ramachandran, 2019; Xiao *et al*, 2024). Interestingly, studies in mammalian systems have shown that the Yme1 ortholog YME1L is itself regulated by mitochondrial lipid composition (MacVicar *et al*, 2019), including PE levels, suggesting that lipid homeostasis and Yme1-family proteases may participate in conserved feedback mechanisms that couple mitochondrial membrane composition to protease activity. In parallel, the MICOS complex organizes crista junctions and maintains associations between the IMM and OMM (Körner *et al*, 2012; Ott *et al*, 2012; Zerbes *et al*, 2012; Aaltonen *et al*, 2016; Cogliati *et al*, 2016). Loss of MICOS may therefore disrupt membrane associations or alter lipid distribution between mitochondrial subcompartments, indirectly promoting OMM remodeling into MDCs. In this framework, Yme1-dependent remodeling of lipid transfer pathways and mitochondrial membrane organization relieves constraints that would otherwise limit the formation of MDCs. Notably, additional Yme1-sensitive proteins were altered in the proteomics dataset, including IMS-localized components of mitochondrial protein import, suggesting that the partly incomplete bypass of Yme1 by combining *ups2Δ* and *MICOSΔ* may reflect contributions from additional downstream effectors.

Our findings further indicate that Yme1-dependent remodeling occurs constitutively but is somehow responsive to metabolic stress. Several Yme1-regulated proteins change in abundance even under steady-state conditions and undergo further remodeling following MDC-inducing treatments such as rapamycin. Consistent with this idea, increasing Yme1 levels through overexpression is sufficient to induce MDC formation in otherwise untreated cells. However, this effect remains dependent on metabolic context, as Yme1 overexpression does not stimulate MDCS when cells are shifted to amino-acid-free media. These observations suggest that metabolic signals gate MDC formation by modulating either Yme1 activity itself or its substrates, or other downstream steps required for compartment formation.

Based on these observations, we propose the model summarized in Figure 5C. In this model, Yme1 acts as a central regulator that promotes MDC formation by progressively removing lipid and organizational constraints that normally stabilize mitochondrial membranes. Under basal conditions, Yme1-mediated turnover of key substrates contributes to ongoing mitochondrial remodeling but is insufficient to trigger widespread MDC formation. During metabolic stress, however, this remodeling is amplified—either through increased Yme1 activity or alterations in its substrates—allowing MDC biogenesis to proceed. In this way, MDC formation emerges as the outcome of coordinated changes in mitochondrial lipid composition, membrane organization, and metabolic state. Together, these findings position Yme1 as a key integrator of mitochondrial proteostasis, membrane organization, and metabolic signaling in the control of MDC formation.

## MATERIALS AND METHODS

### Yeast strains

*Saccharomyces cerevisiae* strains used in this study are described in Table S1 and are derivatives of either S288C (Baker Brachmann *et al*, 1998) or W303 (Matheson *et al*, 2017). All deletion mutants described in this study were generated via sporulation. In a diploid, one allele of the target gene was deleted by PCR mediated gene replacement using the previously described pRS series of vectors (Sikorski & Hieter; Andersen, 2012) and oligo pairs listed in Table S2. Insertion of the selection cassette was verified by colony PCR across the chromosomal integration site. The confirmed heterozygous diploids gene deletions were subsequently sporulated to generate haploid mutants. Strains expressing fluorescently tagged proteins were created by PCR-mediated C-terminal endogenous epitope tagging using standard techniques and oligo pairs listed in Table S2. Plasmid templates for fluorescent epitope tagging were from the pKT series of vectors (Sheff & Thorn, 2004). Correct integrations were confirmed by correctly localized expression of the fluorophore by microscopy. Further, HA-tagged strains were generated by PCR-mediated C-terminal endogenous tagging using pFA6A-3xHA-KanMX (Bähler *et al*, 1998) as the template plasmid and oligonucleotides listed in Table S2, with correct integration confirmed by colony PCR and western blotting. Yeast strains AHY 15346, AHY 15348, and AHY15438 were rendered prototrophic with pHLUM (Mülleder *et al*, 2012) to eliminate complications arising from amino acid auxotrophies and were subsequently used to assess amino acid dependencies on MDC formation.

### Plasmids

Plasmids used in this study are listed in Table S3. pHLUM, a yeast plasmid expressing multiple auxotrophic marker genes from their endogenous promoters, was obtained from Addgene (# 40276) (Mülleder *et al*, 2012). Plasmids for GPD driven expression of *YME1* were generated by Gateway mediated transfer of corresponding ORF (Harvard Institute of Proteomics) from pDONR221 into a pAG306-ccdB chromosome I (Hughes & Gottschling, 2012) using Gateway LR Clonase II enzyme mix (ThermoFisher) according to the manufacturer’s instructions. All insert sequences were verified by Azenta/Genewiz sequencing. The resulting expression plasmid was digested with NotI and integrated into yeast chromosome I (199456-199457).

### Cell culture and media

Yeast cells were grown exponentially for 15–16 h at 30°C (OD₆₀₀= 0.2–0.8) before the start of all experiments. This period of overnight log-phase growth is required to ensure mitochondrial uniformity across the cell population and is critical for MDC biogenesis. Unless indicated, cells were cultured in YPAD medium (1% yeast extract, 2% peptone, 0.005% adenine, 2% glucose). For amino acid dependency experiments, SD medium (low amino acid medium) (0.67% yeast nitrogen base without amino acids, 2% glucose, supplemented with the following nutrients 0.074 g/liter each of adenine, alanine, arginine, asparagine, aspartic acid, cysteine, glutamic acid, glutamine, glycine, histidine, myoinositol, isoleucine, lysine, methionine, phenylalanine, proline, serine, threonine, tryptophan, tyrosine, uracil, valine, 0.369 g/liter of leucine, 0.007 g/liter of para-aminobenzoic acid) or minimal medium i.e. no amino acid medium (0.67% yeast nitrogen base without amino acids, 2% glucose) was used. Where indicated, cells were treated with rapamycin, concanamycin A, or cycloheximide at final concentrations of 200nM, 500nM, or 10ug/mL respectively to induce MDC biogenesis.

### MDC assays

For MDC assays, overnight log-phase cultures were either directly harvested or after a two-hour treatment with dimethyl sulfoxide (DMSO) or the indicated drug. For MDC assays with plasmid-containing cells, overnight log-phase yeast cultures were grown in selective SD medium, back-diluted to an OD₆₀₀ of 0.2–0.4 in YPAD, and treated with DMSO or the indicated MDC inducer for 2 h. Prior to visualization, cells were harvested by centrifugation, washed once with distilled water, resuspended in an imaging buffer (5% wt/vol Glucose, 10 mM HEPES, pH 7.6) and imaged as described below.

### Microscopy and image analysis

Yeast were plated onto a slide at small volumes to allow for the formation of a monolayer. Optical z-sections of live yeast cells were acquired with a Zeiss Axiocam 506 monochromatic camera and 63 x oil immersion objectives (Carl Zeiss, Plan Apochromat, NA 1.4). To obtain super-resolution images a ZEISS LSM800 confocal microscope equipped with an Airyscan detector and 63 x oil immersion objective (Carl Zeiss, Plan Apochromat, NA 1.4) was used. Maximum intensity projected widefield images generated in Zen (Carl Zeiss) were used to quantify the percentage of cells with MDCs. MDCs were identified as Tom70-positive, Tim50-negative structures that were enriched for Tom70 versus the mitochondrial tubule. Unless indicated, maximum-intensity projected images are displayed for all images. All MDC quantifications show the mean ± SE from at least three biological replicates with n≥100 cells per experiment. Representative images were processed using Fiji. Maximum intensity projections of individual channels were generated, and fluorescence intensities were minimally adjusted for visualization to enable comparison across channels. Line-scan analyses were performed on non-adjusted single z-sections.

### Protein preparation and immunoblotting

Yeast cultures were grown overnight as described and treated with either DMSO or Rap. Post treatment, 2 OD₆₀₀ units of cells were collected, washed twice with distilled water and incubated in 0.1M NaOH for 5 min at RT. The pellet was reisolated at 14,000 × g for 10 min at 4°C and resuspended in lysis buffer (10 mM Tris, pH 6.8; 100 mM NaCl; 1 mM EDTA; 1 mM EGTA; 1% SDS; cOmplete™ protease inhibitor cocktail, Millipore Sigma) for 5 min at 95°C. Subsequently, samples were denatured in Laemmli buffer (63 mM Tris pH 6.8, 2% SDS, 10% glycerol, 1 mg/ml bromophenol blue, 1% β-mercaptoethanol) for 5 min at 95°C. The prepared protein samples were resolved on a 4–12% SDS–polyacrylamide gradient gel and transferred onto nitrocellulose membranes using a semi-dry transfer system. To prevent nonspecific antibody binding, membranes were blocked in Tris-buffered saline with 0.05% Tween-20 (TBST) containing 10% nonfat dry milk for 40 min at RT, followed by overnight incubation with indicated primary antibodies at 4°C. Membranes were washed four times with TBST and incubated with secondary antibody (donkey anti-mouse HRP-conjugated, 1:4000 in TBST containing 10% nonfat dry milk; Sigma-Aldrich) for 1 h at room temperature. Following incubation, membranes were washed twice with TBST and twice with TBS. Blots were developed using enhanced chemiluminescence substrate (Thermo Fisher Scientific), and signals were detected with a Bio-Rad ChemiDoc MP system. Images were exported as TIFF files and cropped in Adobe Illustrator.

### Analysis of protein levels using FIJI

Raw TIFF images were imported into FIJI for densitometric analysis. A standardized rectangular region of interest was applied to each band to obtain the integrated density (AU). To correct for background, four measurements were taken from blank regions of each blot and averaged. Each band intensity was normalized first to its respective loading control within the same lane and then to the corresponding wild-type untreated sample. Quantification was performed using three independent replicates per condition. Data were plotted in GraphPad Prism, and p-values were calculated using two-way ANOVA.

### Quantification and statistical analysis

No randomization or blinding was performed, as all experiments were conducted using defined laboratory reagents and yeast strains with known genotypes. For all experiments, biological replicates are represented by “n,” with at least three replicates per experiment. All statistical analyses were conducted in GraphPad Prism, with the specific tests and measures of dispersion and precision reported in the figure legends.

### Mitochondrial proteomics Isolation of yeast mitochondria

Yeast were grown overnight to log-phase (OD₆₀₀= 0.4–0.8) and treated with either DMSO or Rap for 2 h. Subsequently, cells were isolated by centrifugation, washed with distilled water, and the pellet weight was recorded. The pellets were resuspended in dithiothreitol (DTT) buffer (0.1 M Tris, 10 mM DTT, pH 9.4) at 2mL per gram of the pellet and incubated for 20 min at 30°C under constant rotation. After reisolation by centrifugation, DTT treated cells were washed once in 1.2M sorbitol and resuspended in sorbitol phosphate buffer (6.7 mL per gram of pellet) containing 2 mg of lyticase per gram of pellet. Cells were incubated for 30–45 min at 30°C with constant rotation to digest the cell wall and generate spheroplasts. Following lyticase treatment, spheroplasts were isolated by centrifugation and lysed by mechanical disruption in homogenization buffer (0.6 M sorbitol, 10 mM Tris, pH 7.4, 1 mM EDTA, pH 8.0 adjusted with KOH, 0.2% BSA, 1 mM PMSF) at 13.3 mL per gram of pellet at 4°C. Cell debris was removed from the homogenate twice by centrifugation at 2,000 × g for 5 min each at 4°C and mitochondria were pelleted at 17,500 × g for 12 min at 4°C. The mitochondrial pellet was resuspended in SEM buffer (250 mM sucrose, 1 mM EDTA, pH 8.0 adjusted with KOH, 10 mM 3-(N-morpholino)-prop-ansulfonic acid, pH 7.2), and reisolated by differential centrifugation as described above, and subsequently resuspended in SEM buffer. The protein concentration of mitochondria was determined by Bicinchoninic Acid (BCA) Assay. For proteomics analysis, 100 μg of mitochondria (by protein) were pelleted, shock-frozen in liquid nitrogen, and stored at −80°C.

### Mass spectrometry sample preparation

Samples for protein analysis were prepared essentially as previously described (Navarrete-Perea *et al*, 2018; Li *et al*, 2021). Proteomes were extracted using a buffer containing 200 mM EPPS pH 8.5, 8M urea, 0.1% SDS and protease inhibitors. Following lysis, 25 µg of each proteome was reduced with 5 mM TCEP. Cysteine residues were alkylated using 10 mM iodoacetimide for 20 minutes at RT in the dark. Excess iodoacetimide was quenched with 10 mM DTT. A buffer exchange was carried out using a modified SP3 protocol (Hughes *et al*, 2019). Briefly, ∼250 µg of Cytiva SpeedBead Magnetic Carboxylate Modified Particles (65152105050250 and 4515210505250), mixed at a 1:1 ratio, were added to each sample. 100% ethanol was added to each sample to achieve a final ethanol concentration of at least 50%. Samples were incubated with gentle shaking for 15 minutes. Samples were washed three times with 80% ethanol. Protein was eluted from SP3 beads using 200 mM EPPS pH 8.5 containing Lys-C (Wako, 129-02541). Samples were digested overnight at room temperature with vigorous shaking. The next morning trypsin was added to each sample and further incubated for 6 hours at 37° C. Acetonitrile was added to each sample to achieve a final concentration of ∼33%. Each sample was labelled, in the presence of SP3 beads, with ∼62.5 µg of TMTPro reagents (ThermoFisher Scientific). Following confirmation of satisfactory labelling (>97%), excess TMT was quenched by addition of hydroxylamine to a final concentration of 0.3%. The full volume from each sample was pooled and acetonitrile was removed by vacuum centrifugation for 1 hour. The pooled sample was acidified and peptides were de-salted using a Sep-Pak 50mg tC18 cartridge (Waters). Peptides were eluted in 70% acetonitrile, 1% formic acid and dried by vacuum centrifugation.

### Basic pH reversed-phase separation (BPRP)

TMT labeled peptides were solubilized in 5% acetonitrile/10 mM ammonium bicarbonate, pH 8.0 and ∼300 µg of TMT labeled peptides were separated by an Agilent 300 Extend C18 column (3.5 mm particles, 4.6 mm ID and 250 mm in length). An Agilent 1260 binary pump coupled with a photodiode array (PDA) detector (Thermo Scientific) was used to separate the peptides. A 45 minute linear gradient from 10% to 40% acetonitrile in 10 mM ammonium bicarbonate pH 8.0 (flow rate of 0.6 mL/min) separated the peptide mixtures into a total of 96 fractions (36 seconds). A total of 96 Fractions were consolidated into 24 samples in a checkerboard fashion and vacuum dried to completion. Each sample was desalted via Stage Tips and re-dissolved in 5% formic acid/ 5% acetonitrile for LC-MS3 analysis.

### Liquid chromatography and tandem mass spectrometry

Mass spectrometric data were collected on an Orbitrap Eclipse mass spectrometer coupled to a Proxeon NanoLC-1000 UHPLC (Thermo Fisher Scientific). The 100 µm capillary column was packed in-house with 35 cm of Accucore 150 resin (2.6 μm, 150Å; ThermoFisher Scientific). Data were acquired for 120 min per run. A FAIMS device was enabled during data collection and compensation voltages were set at -40V, -60V, and -80V (Schweppe *et al*, 2019). MS1 scans were collected in the Orbitrap (resolution – 60,000; scan range – 400-1600 Th; automatic gain control (AGC) – 4x10^5^, maximum ion injection time – automatic). MS2 scans were collected in the Orbitrap following higher-energy collision dissociation (HCD; resolution – 50,000; AGC – 1.25x10^5^; normalized collision energy – 36; isolation window – 0.5 Th; maximum ion injection time – 86 ms.

### Mass spectrometry data analysis

Database searching included all entries from the Saccharomyces cerevisiae UniProt Database (downloaded in June 2023). The database was concatenated with one composed of all protein sequences for that database in the reversed order (Elias JE *et al*, 2007). Raw files were converted to mzXML, and monoisotopic peaks were re-assigned using Monocle (Rad *et al*, 2021). Searches were performed with Comet (Eng *et al*, 2013) using a 50-ppm precursor ion tolerance and fragment bin tolerance of 0.02. TMTpro labels on lysine residues and peptide N-termini (+304.207 Da), as well as carbamidomethylation of cysteine residues (+57.021 Da) were set as static modifications, while oxidation of methionine residues (+15.995 Da) was set as a variable modification. Peptide-spectrum matches (PSMs) were adjusted to a 1% false discovery rate (FDR) using a linear discriminant after which proteins were assembled further to a final protein-level FDR of 1% analysis (Huttlin *et al*, 2010). TMT reporter ion intensities were measured using a 0.003 Da window around the theoretical m/z for each reporter ion. Proteins were quantified by summing reporter ion counts across all matching PSMs. More specifically, reporter ion intensities were adjusted to correct for the isotopic impurities of the different TMTpro reagents according to manufacturer specifications. Peptides were filtered to exclude those with a summed signal-to-noise (SN) < 180 across all TMT channels and < 0.5 precursor isolation specificity. The signal-to-noise (S/N) measurements of peptides assigned to each protein were summed (for a given protein). The mass spectrometry proteomics data have been deposited to the ProteomeXchange Consortium via the PRIDE (Perez-Riverol *et al*, 2025) partner repository with the dataset identifier PXD075997. Table S4 provides a summary of peptide counts and signal-to-noise (S/N) ratios across all identified proteins in all samples.

### Statistical analysis and data visualization

Proteomics data were log_2_-transformed prior to analysis. Principal component analysis (PCA) was performed using the NIPALS algorithm (Wright K, 2025) to evaluate sample clustering. Boxplots with color and pattern fills were used to compare protein expression across treatment and genotype groups, respectively. Differences in protein abundance between groups were assessed using Welch’s two-sample t-tests. Proteins were considered differentially expressed if they exhibited at least a 1.5-fold change in either direction and Storey’s q-value ≤ 0.05 (Storey & Tibshirani, 2003). Volcano plots were generated to visualize fold changes and significance. Z-score normalized values of selected proteins were visualized in heatmaps with hierarchical clustering using the ComplexHeatmap R package (Gu *et al*, 2016; Gu, 2022). Protein annotations were retrieved from UniProt (The UniProt Consortium *et al*, 2025) on 2025-02-12 to obtain Gene Ontology (GO) terms. Table S5 details the Gene Ontology (GO) annotations associated with the proteins identified in Table S4. Pathway enrichment analysis was performed using the gprofiler2 (Kolberg *et al*, 2020) R package (organism = "scerevisiae"), which queries multiple databases by default. For visualization and interpretation, enrichment results were filtered to the GO:Biological Process (GO:BP) category. Table S6 summarizes the corresponding functional enrichment results generated using g:Profiler.

### Whole-cell lipidomics sample preparation

Yeast cells were grown overnight as described above and treated with the indicated inducers. Following treatment, 5 OD₆₀₀ units of cells were harvested, washed twice with dH₂O, and flash-frozen. Extraction of lipids was carried out using a biphasic solvent system of cold methanol, methyl *tert*-butyl ether (MTBE), and PBS (Matyash *et al*, 2008) with some modifications. In a randomized sequence, to each sample was added 230 µL MeOH with internal standards. Samples were homogenized for 30 seconds, transferred to microcentrifuge tubes (polypropylene 1.7 mL, VWR, USA) containing 750 µL MTBE, and then incubated on ice with occasional vortexing for 1 hr. Following incubation, samples were centrifuged at 15,000 x g for 10 minutes at 4°C. The organic (upper) layer was collected, and the aqueous (lower) layer was re-extracted with 1 mL of 10:3:2.5 (*v/v/v*) MTBE/MeOH/dd-H2O, briefly vortexed, incubated at RT, and centrifuged at 15,000 x g for 10 minutes at 4°C. Upper phases were combined and evaporated to dryness under speedvac. Lipid extracts were reconstituted in 250 µL of 4:1:1 (v/v/v) IPA/ACN/water and transferred to LC-MS vials for analysis. Concurrently, a process blank sample was prepared and pooled quality control (QC) samples were prepared by taking equal volumes from each sample after final resuspension.

### LC-MS analysis (QTOF)

Lipid extracts were separated on an Acquity UPLC CSH C18 column (2.1 x 100 mm; 1.7 µm) coupled to an Acquity UPLC CSH C18 VanGuard precolumn (5 × 2.1 mm; 1.7 µm) (Waters, Milford, MA) maintained at 65°C connected to an Agilent HiP 1290 Sampler, Agilent 1290 Infinity pump, and Agilent 6545 Accurate Mass Q-TOF dual AJS-ESI mass spectrometer (Agilent Technologies, Santa Clara, CA). Samples were analyzed in a randomized order in both positive and negative ionization modes in separate experiments acquired with the scan range m/z 100 – 1700. For positive mode, the source gas temperature was set to 225°C, with a drying gas flow of 11 L/min, nebulizer pressure of 40 psig, sheath gas temp of 350°C and sheath gas flow of 11 L/min. VCap voltage is set at 3500 V, nozzle voltage 500V, fragmentor at 110 V, skimmer at 85 V and octopole RF peak at 750 V. For negative mode, the source gas temperature was set to 300°C, with a drying gas flow of 11 L/min, a nebulizer pressure of 30 psig, sheath gas temp of 350°C and sheath gas flow 11 L/min. VCap voltage was set at 3500 V, nozzle voltage 75 V, fragmentor at 175 V, skimmer at 75 V and octopole RF peak at 750 V. Mobile phase A consisted of ACN:H_2_O (60:40, *v/v*) in 10 mM ammonium formate and 0.1% formic acid, and mobile phase B consisted of IPA:ACN:H_2_O (90:9:1, *v/v/v*) in 10 mM ammonium formate and 0.1% formic acid. For negative mode analysis the modifiers were changed to 10 mM ammonium acetate. The chromatography gradient for both positive and negative modes started at 15% mobile phase B then increased to 30% B over 2.4 min, it then increased to 48% B from 2.4 – 3.0 min, then increased to 82% B from 3 – 13.2 min, then increased to 99% B from 13.2 – 13.8 min where it’s held until 16.7 min and then returned to the initial conditions and equilibrated for 5 min. Flow was 0.4 mL/min throughout, with injection volumes of 1 µL for positive and 10 µL negative mode.

Tandem mass spectrometry was conducted using iterative exclusion, the same LC gradient at collision energies of 20 V and 27.5 V in positive and negative modes, respectively.

### LC-MS data processing

For data processing, Agilent MassHunter (MH) Workstation and software packages MH Qualitative and MH Quantitative were used. The pooled QC (n=8) and process blank (n=4) were injected throughout the sample queue to ensure the reliability of acquired lipidomics data. For lipid annotation, accurate mass and MS/MS matching was used with the Agilent Lipid Annotator library and LipidMatch (Koelmel *et al*, 2017). Results from the positive and negative ionization modes from Lipid Annotator were merged based on the class of lipid identified. Data exported from MH Quantitative was evaluated using Excel where initial lipid targets are parsed based on the following criteria. Only lipids with relative standard deviations (RSD) less than 30% in QC samples are used for data analysis. Additionally, only lipids with background AUC counts in process blanks that are less than 30% of QC are used for data analysis. The parsed excel data tables are normalized based on the ratio to class-specific internal standards, then to tissue mass and sum prior to statistical analysis.

### Statistical analysis and data visualization

Four biological replicates were used for each lipidomics experiment and analyzed within the same LC-MS run. Statistical analysis was performed in GraphPad Prism using two-way ANOVA with Holm–Sidak post hoc tests, as indicated in the figure legends. All data were included except those that did not meet QC criteria; if a replicate failed QC, the entire biological replicate was excluded from further analysis. For each denoted phospholipid, all corresponding lipid species were considered, and values were normalized to OD of whole-cell lysates. Table S7 summarizes cumulative lipid levels across samples, including PE abundance and the corresponding percentage of PE relative to total lipid content.

## ONLINE SUPPLEMENTAL MATERIAL

Figure S1 shows that secondary effects related to loss of Yme1 do not impact MDC formation. Figure S2 depicts quality control analyses of the TMT-based mitoproteomics dataset including data normalization (box plots) and principal component analysis (PCA). Figure S3 shows MDC assay profiling of Yme1-regulated factors in the MDC pathway. Figure S4 highlights the role of the MICOS complex in MDC biogenesis, including the effects of individual MICOS mutants and their genetic interactions with Yme1 and Ups2. Table S1 lists the yeast strains used in this study. Table S2 contains the oligonucleotides used in this study. Table S3 includes bacterial strains, chemicals, antibodies, plasmids, and software used in this study. Table S4 contains the TMT-based quantitative proteomics dataset, containing the peptide counts and scaled signal-to-noise (S/N) values for all identified proteins across all samples. Table S5 lists the Gene Ontology (GO) annotations associated with the identified proteins listed in Table S4. Table S6 contains the functional enrichment analysis performed using g:Profiler. Table S7 summarizes total lipid levels, including the relative and/or absolute abundance of PE levels measured across the indicated conditions.

## DATA AVAILABILITY

All reagents used in this study are available upon request. All other data reported in this paper will be shared by the lead contact upon request. This paper does not report the original code. Any additional information required to reanalyze the data reported in this paper is available from the lead contact upon request.

## ACKNOWLEDGEMENTS

We thank past and present members of the A.L. Hughes group for their discussion and manuscript comments. We thank members of Janet. M. Shaw laboratory for providing reagents, mitochondrial antibodies, and support early in the project. Proteomics analysis was conducted at the Thermo Fisher Scientific Center for Multiplexed Proteomics at Harvard Medical School. We thank Jonathan Van Vranken, PhD for performing the TMT-based proteomics experiments. We also thank Paul Stewart, PhD for his assistance with subsequent data analysis and visualizations. Lipidomics analysis was performed at the Metabolomics Core Facility at the University of Utah.

We sincerely thank John Alan Maschek, PhD for his expertise and assistance with the lipidomic analyses. Research was supported by National Institutes of Health grants GM119694 and AG061376 to A.L. Hughes and a University of Utah Graduate Research Fellowship to S.S. Balasubramaniam. J.F. is supported by NIH grant GM137894. The content is solely the responsibility of the authors and does not necessarily represent the official views of the NIH.

## AUTHOR CONTRIBUTIONS

Conceptualization, S.S. Balasubramaniam, J.R. Friedman, and A.L. Hughes; methodology, S.S. Balasubramaniam; formal analysis, S.S. Balasubramaniam; investigation, S.S. Balasubramaniam, A.E. Curtis; writing – original draft, S.S. Balasubramaniam; writing – review and editing, S.S. Balasubramaniam, J.R. Friedmann and A.L. Hughes; visualization, S.S. Balasubramaniam and A.L. Hughes; supervision, A.L. Hughes; funding acquisition, S.S. Balasubramaniam, J.R. Friedman, and A.L. Hughes.

## CONTACT FOR REAGENT AND RESOURCE SHARING

Further information and requests for resources and reagents should be directed to and will be fulfilled by the Lead Contact, Adam Hughes. All unique/stable reagents generated in this study are available from the Lead Contact without restrictions.

---

**Figure S1:**
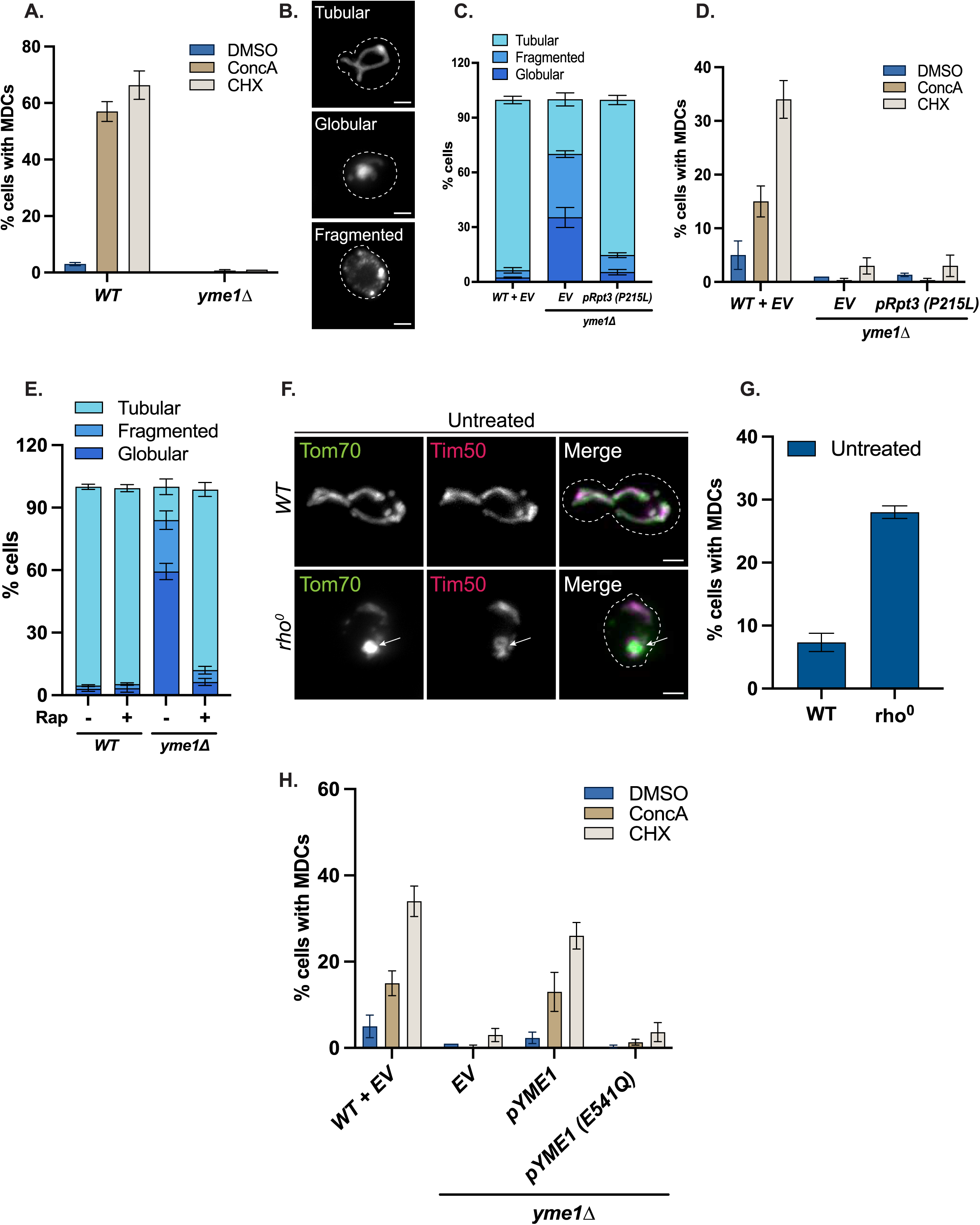
Secondary effects related to loss of Yme1 do not impact MDC formation (Related to Figure 1) (A) Quantification of MDC formation in wild-type and *yme1Δ* cells treated with DMSO, ConcA, or CHX for 2 h. Error bars represent the mean ± SE of three replicates, *n* ≥ 100 cells per replicate. (B) Representative widefield images of distinct yeast mitochondrial morphologies. Scale bar = 2 µm. (C) Quantification of mitochondrial morphology in wild-type and *yme1Δ* cells expressing either *EV* or *pRS413-Rpt3 (P215L)*. Error bars represent the mean ± SE of three replicates, *n* ≥ 100 cells per replicate. (D) Quantification of MDC formation in wild-type and *yme1Δ* cells expressing either *EV* or *pRS413-Rpt3 (P215L)* treated with DMSO, ConcA, or CHX for 2 h. Error bars represent the mean ± SE of three replicates, *n* ≥ 100 cells per replicate. (E) Quantification of mitochondrial morphology in wild-type and *yme1Δ* cells treated with DMSO or Rap for 2 h. Error bars represent the mean ± SE of three replicates, *n* ≥ 100 cells per replicate. (F) Widefield images of wild-type and *rho^0^* cells expressing Tom70-GFP and Tim50-mCherry. White arrows denote MDCs. White arrows denote MDCs. Scale bar = 2 µm. (G) Quantification of (F) showing the percentage of cells with MDCs. Error bars represent the mean ± SE of three replicates, *n* ≥ 100 cells per replicate. (H) Quantification of MDC formation in wild-type and *yme1Δ* cells expressing either *EV*, pRS413-Yme1, or pRS413-Yme1 (E541Q) treated with DMSO, ConcA, or CHX for 2 h. Error bars represent the mean ± SE of three replicates, *n* ≥ 100 cells per replicate.

**Figure S2:**
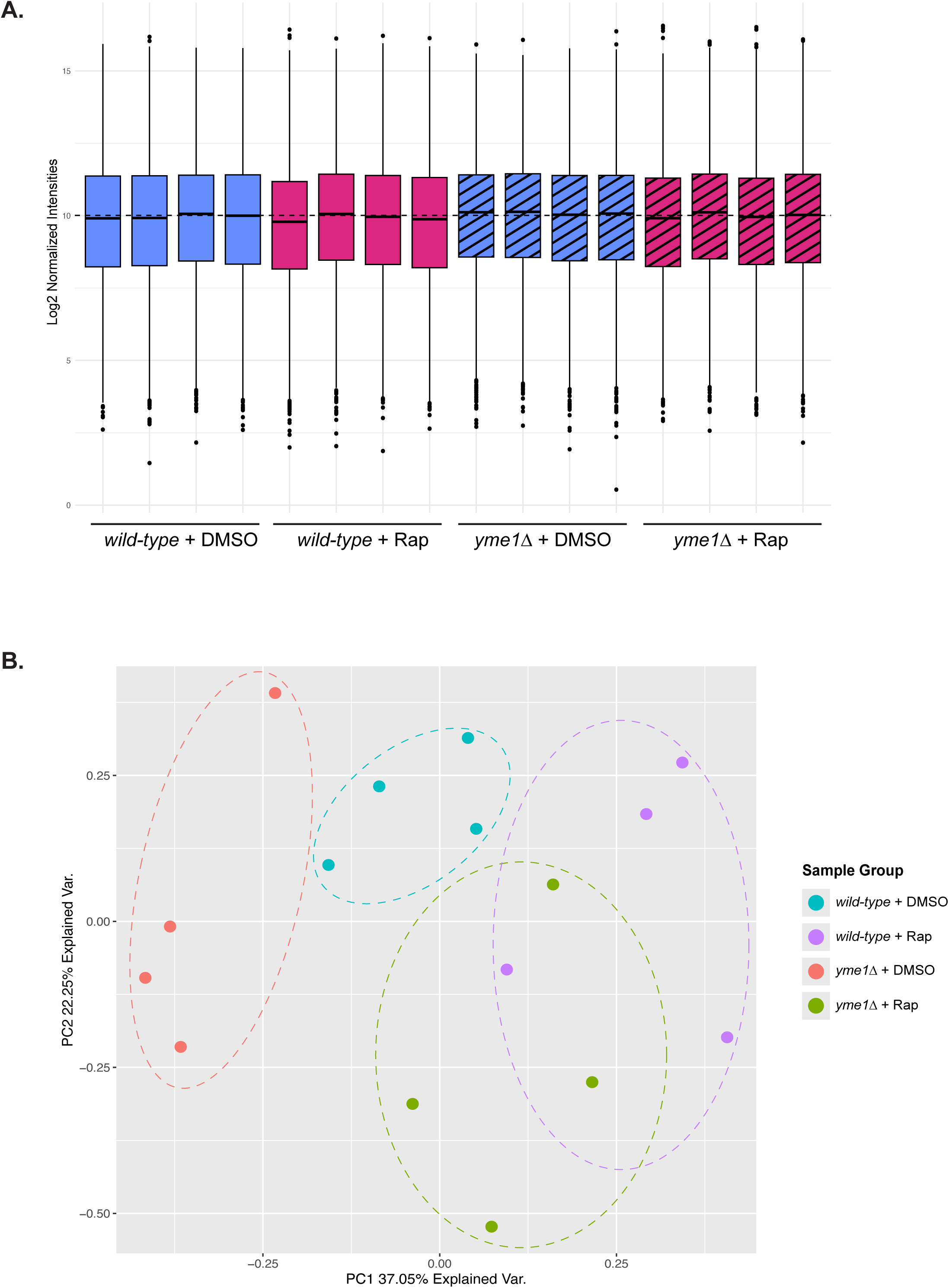
Mitoproteomics data quality control (Related to Figure 2) (A) Box plot showing log2-normalized protein intensities between wild-type and *yme1Δ* cells treated with DMSO or Rap for 2 h (n = 4). Comparable distributions across conditions confirm successful data normalization. (B) Principal component analysis of proteome data from wild-type and *yme1Δ* cells treated with DMSO or Rap for 2 h (n = 4).

**Figure S3:**
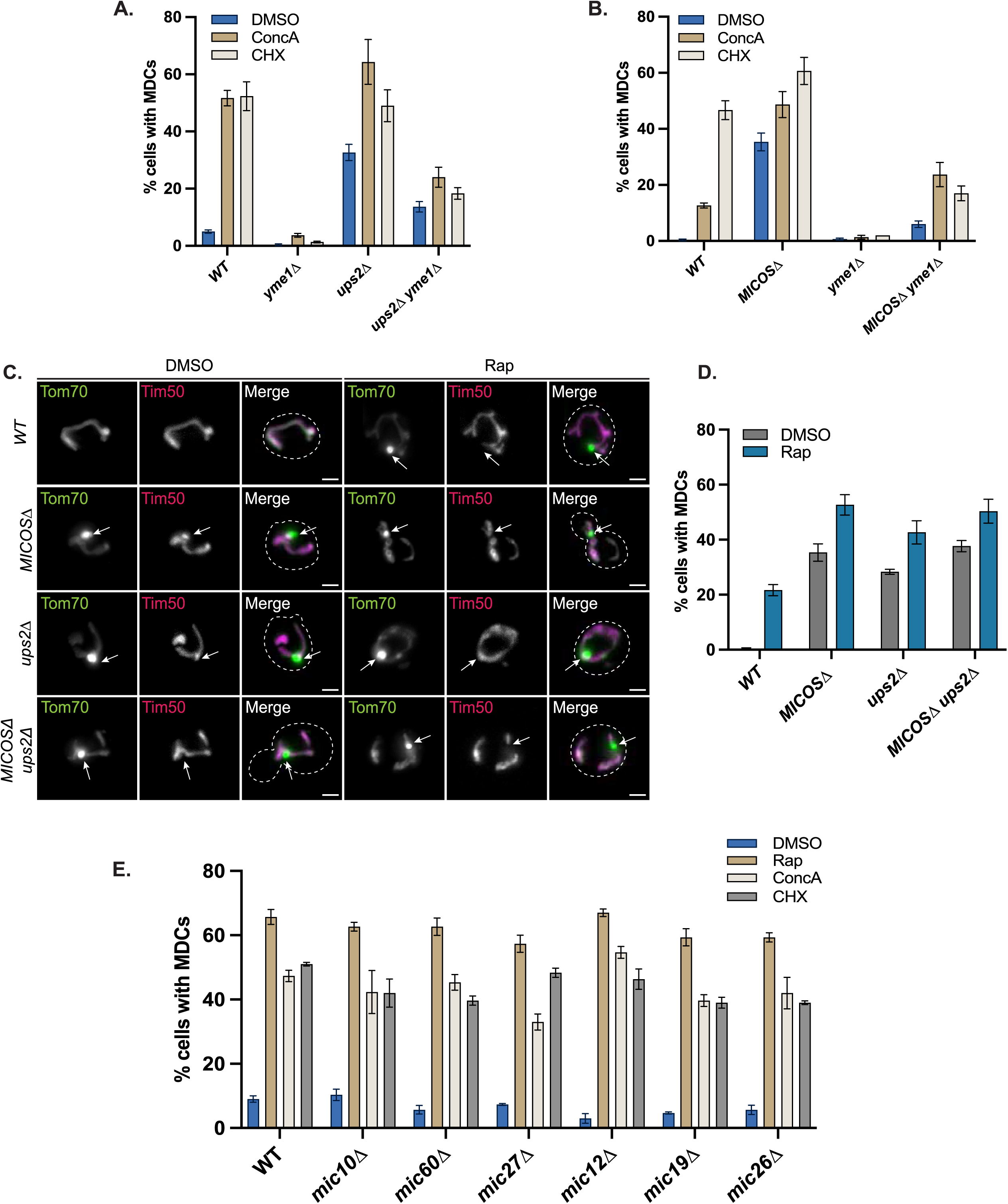
MDC assay profiling of Yme1-regulated factors (Related to Figures 3 & 4) (A) Quantification of MDC formation in wild-type and indicated mutants treated with DMSO, ConcA, or CHX for 2 h. Error bars represent the mean ± SE of three replicates, *n* ≥ 100 cells per replicate. (B) Quantification of MDC formation in wild-type and indicated mutants treated with DMSO, ConcA, or CHX for 2 h. Error bars represent the mean ± SE of three replicates, *n* ≥ 100 cells per replicate. (C) Widefield images of wild-type and indicated mutants expressing Tom70-GFP and Tim50-mCherry treated with DMSO or Rap for 2 h. White arrows denote MDCs. Scale bar = 2 µm. (D) Quantification of (C) showing the percentage of cells with MDCs. Error bars represent the mean ± SE of three replicates, *n* ≥ 100 cells per replicate. (E) Quantification of MDC formation in wild-type and indicated mutants treated with DMSO, Rap, ConcA, or CHX for 2 h. Error bars represent the mean ± SE of three replicates, *n* ≥ 100 cells per replicate.

**Figure S4:**
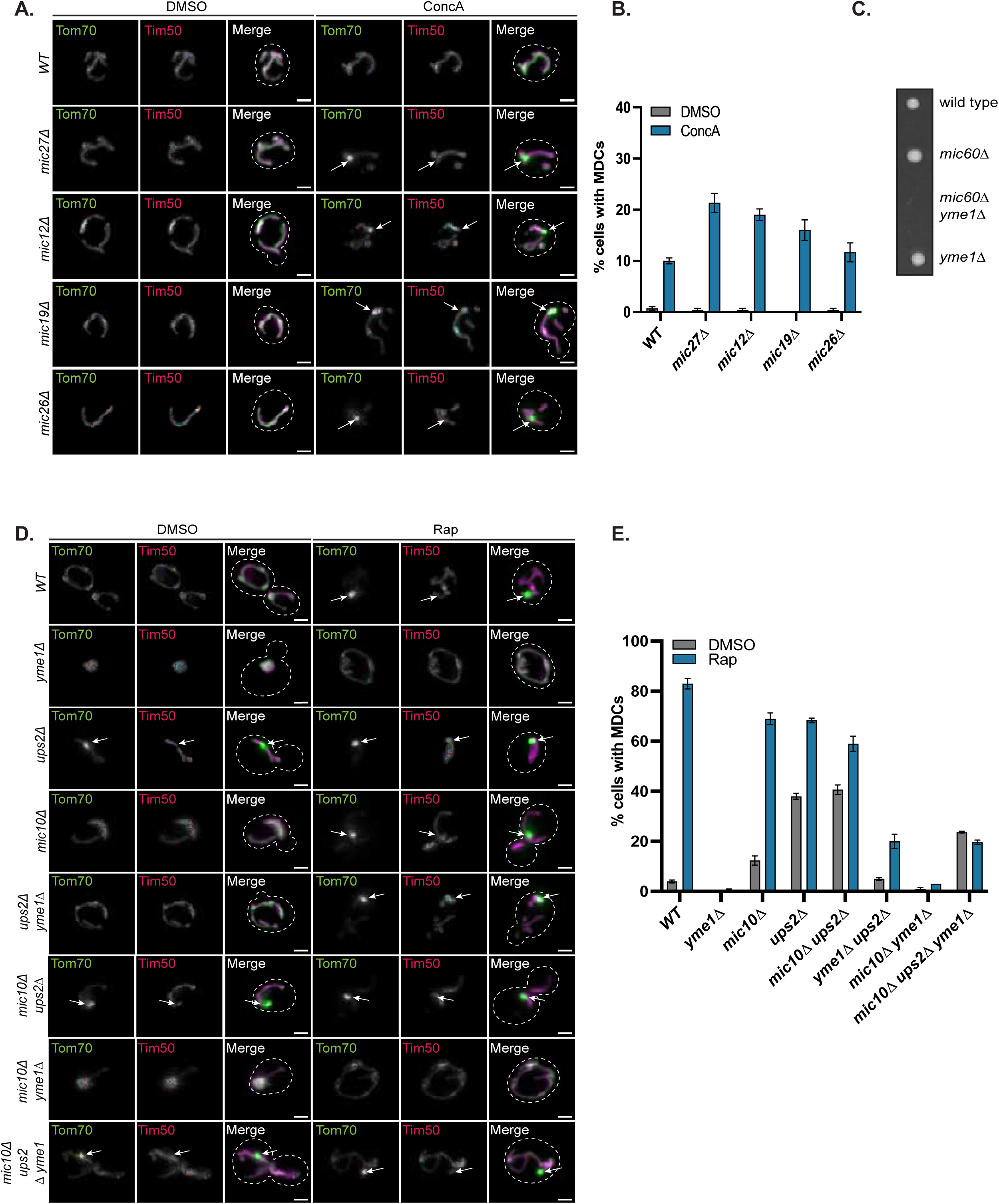
Role of the MICOS Complex in MDC formation: Effects of individual mutants and genetic interactions with Yme1, and Ups2 (Related to Figure 4) (A) Widefield images of wild-type and indicated mutants expressing Tom70-GFP and Tim50-mCherry, grown in media containing low amino acids and treated with DMSO or ConcA for 2 h. White arrows denote MDCs. Scale bar = 2 µm. (B) Quantification of (A) showing the percentage of cells with MDCs. Error bars represent the mean ± SE of three replicates, *n* ≥ 100 cells per replicate. (C) Tetrad dissection of *mic60Δ/+ yme1Δ/+* diploid cells demonstrating synthetic lethality between *MIC60* and *YME1*. (D) Widefield images of wild-type and indicated mutants expressing Tom70-GFP and Tim50-mCherry treated with DMSO or Rap for 2 h. White arrows denote MDCs. Scale bar = 2 µm. (E) Quantification of (D) showing the percentage of cells with MDCs. Error bars represent the mean ± SE of three replicates, *n* ≥ 100 cells per replicate.

